# Breaking the barrier of human-annotated training data for machine-learning-aided plant research using aerial imagery

**DOI:** 10.1101/2025.04.02.646808

**Authors:** Sebastian Varela, Xuying Zheng, Joyce Njuguna, Erik Sacks, Dylan Allen, Jeremy Ruhter, Andrew D.B. Leakey

**Affiliations:** Center for Advanced Bioenergy and Bioproducts Innovation; University of Illinois at Urbana Champaign; Urbana, IL 61801, USA; Independent Researcher; Canelones, 15800, Uruguay; Department of Crop Sciences, University of Illinois at Urbana Champaign; Urbana, IL 61801, USA; Institute for Genomic Biology, University of Illinois at Urbana Champaign; Urbana, IL 61801, USA; Department of Plant Biology, University of Illinois at Urbana Champaign; Urbana, IL 61801, USA; Center for Digital Agriculture, University of Illinois at Urbana Champaign; Urbana, IL 61801, USA

**Keywords:** Plant Phenotyping, Generative and Adversarial Learning, Uncrewed aerial vehicle (UAV), Uncrewed aerial system (UAS), drone

## Abstract

Machine learning (ML) can accelerate biological research. However, the adoption of such tools to facilitate phenotyping based on sensor data has been limited by: (1) the need for a large amount of human-annotated training data for each context in which the tool is used; and (2) phenotypes varying across contexts defined in terms of genetics and environment. This is a major bottleneck because acquiring training data is generally costly and time-consuming. This study demonstrates how a ML approach can address these challenges by minimizing the amount of human supervision needed for tool building. A case study comparing ML approaches that examine images collected by an uncrewed aerial vehicle was performed to determine the presence of panicles (i.e., “heading”) across thousands of field plots containing genetically diverse breeding populations of two *Miscanthus* species. Automated analysis of aerial imagery enabled the identification of heading approximately nine times faster than in-field visual inspection by humans. Leveraging an Efficiently Supervised Generative and Adversarial Network (ESGAN) learning strategy reduced the requirement for human-annotated data by one to two orders of magnitude compared to traditional, fully supervised learning approaches. The ESGAN model learned the salient features of the dataset by using thousands of unlabeled images to inform the discriminative ability of a classifier so that it required minimal human-labeled training data. This method can accelerate the phenotyping of heading date as a measure of flowering time in Miscanthus across diverse contexts (e.g., in multi-state trials) and opens avenues to promote the broad adoption of ML tools.

**One-sentence summary:** Machine learning approach accelerates plant phenotyping by reducing the need for large amounts of human-annotated data, enabling faster and more efficient detection of a plant trait using aerial imagery

The author responsible for distribution of materials integral to the findings presented in this article in accordance with the policy described in the Instructions for Authors (https://academic.oup.com/plphys/pages/General-Instructions) is Sebastian Varela.

## Introduction

Artificial intelligence (AI) and machine learning (ML) present enormous opportunities for accelerating scientific discovery, especially in biological research where large-scale, complex problems are commonplace (Burke and Lobell, 2017; Xie et al., 2021; Varela et al., 2022b; Wang et al., 2023). However, advanced AI/ML methods require substantial amounts of annotated data for training purposes and are highly context dependent i.e. they do not perform reliably in contexts beyond that covered by the training data (Paullada et al., 2021; Wang et al., 2022). Biological research is an especially challenging use case for image classification problems because the appearance and function (i.e. phenotype) of an organism is the result of complex interactions between genotype, natural environment and human intervention (Hayes et al., 2023). For example, the performance of a particular crop depends on variation in genotype, growing conditions and management practices (Zhao et al., 2022; Cooper et al., 2023). This contrasts with everyday objects, which can take a variety of forms, but are fixed in time and space such that a given object (e.g. a teapot) will not change shape, texture or color depending on the location and time at which it is imaged under standardized conditions. The potential for AI- and ML-enabled approaches to be applied to biological research has been demonstrated across many scales from cells to organs, organisms, communities and ecosystems. However, this high contextual diversity means existing AI/ML tools will need to be retrained – at considerable cost and effort - for each new biological context in which they are to be used (Moen et al., 2019; Hesami et al., 2022), limiting their adoption. The increasing spatial, temporal, and spectral resolution of sensors, along with cloud computational processing, are all enhancing our ability to supply more sensor data for phenotyping (Lin et al., 2023). However, the greater spatial and temporal resolution of sensor data will only be fully exploited if it is matched by greater resolution in the human annotated data used for training ML tools that find associations between the two datasets. Existing methods to minimize human effort in the production of annotated data include data augmentation (Shorten and Khoshgoftaar, 2019), pseudo-labeling, and label propagation (Iscen et al., 2019; Gan et al., 2022). Meanwhile, active learning (Nagasubramanian et al., 2021), transfer learning (TL) (Tran et al., 2019), and semi-supervised learning (van Engelen and Hoos, 2020) can reduce the need for annotated data in the training process. Recent research efforts to overcome these challenges include developing AI techniques that more rapidly generalize from few examples (Parnami and Lee, 2022). However, while each of these approaches can deliver valuable benefits, the necessity for further innovation to address the intertwined issues of training effort and context dependency is clear (Alzubaidi et al., 2021; Sapoval et al., 2022; Ahmed et al., 2023).

This study proposes a generative-adversarial learning strategy as an alternative to traditional supervised learning and transfer learning, aiming to minimize the human-based supervision required for a computer vision tool. This involves exploiting the ability of a generative-adversarial network (GAN) to learn the salient features of the data from unlabeled images captured with an aerial platform. It is anticipated that allowing the model to learn the underlying latent space representation in the data can be leveraged to enhance the model’s discriminative ability in a classification task with minimal human assistance. A key feature of this approach is using a co-informative learning strategy between the unsupervised and supervised classifiers within the GAN. This is intended to allow learning of the salient features of the large, unlabeled image set to be complemented by use of a smaller pool of labelled images to efficiently achieve the classification task. We describe this approach as an Efficiently Supervised GAN (ESGAN).

A case study of the proposed approach is performed by classifying thousands of diverse, field-grown *Miscanthus* genotypes as having produced panicles, or not, on a given date in a time course of imagery collected by an uncrewed aerial vehicle (UAV, or uncrewed aerial system, or drone). Biomass and valuable chemical compounds from dedicated bioenergy crops are expected to play a central role in the provision of more sustainable energy and bioproducts (Somerville et al., 2010; Martinez-Feria et al., 2022; Eckardt et al., 2023). *Miscanthus sacchariflorus* and *M. sinensis* are crossed to produce very productive, sterile hybrids (Heaton et al., 2008). Flowering time is a key trait influencing productivity and adaptation of Miscanthus to different growing regions (Jensen et al., 2011; Clifton-Brown et al., 2019). Flowering time in Miscanthus, like many other grass crops, can be assessed in terms of “heading date”, i.e. when panicles are outwardly visible in 50% of the culms that reach the top of the canopy (Li et al., 2006; Crowell et al., 2016; Clark et al., 2019; Wu et al., 2019). But, repetitive visual inspections of thousands of individuals grown in extensive field trials is very labor intensive (Clark et al., 2019). Repeated assessment of a crop trial to assess when in a seasonal time course a panicle is first observed then allows estimation of heading date. Increasing the frequency with which the crop is assessed increases the precision of heading date estimates, but also increases labor and has motivated development of ML-enabled remote sensing tools to identify reproductive organs and to assess if plants have reached developmental milestones (Wu et al., 2019; Liu et al., 2020; Fan et al., 2022). In Miscanthus, a fully-supervised 3D-convolutional neural network assessed heading date from UAV images (Varela et al., 2022b). However, the common challenges of context dependency, demand for substantial training data, and limits to generalization ability remain if such tools are to be widely adopted (Rasmussen et al., 2022). Therefore, this is just one of many phenotypes for which reducing the demand for manual supervision of ML tools will be valuable.

This study tests the ability of ESGAN to classify aerial images of individual plants of *M sacchariflorus* and *M. sinensis* on the basis of panicles being visible or not i.e. the most repeated and labor-demanding step in heading date determination. The performance of ESGAN is compared to various popular algorithms based on the fully-supervised learning (FSL) paradigm and traditional TL with varying degrees of complexity, including: K-Nearest Neighbor (KNN), Random Forest (RF), custom Convolutional Neural Network (CNN), and ResNet-50. This analysis was repeated as the number of annotated images provided to train a given model, was reduced from 3137 (100%) to 32 (1%), while simultaneously providing ESGAN with access to the complete set of unannotated images (i.e., n=3137). The objective is to understand the trade-offs between predictive ability and the level of dependence on manual annotation for each of the algorithms. In addition, we test how ESGAN exploits its unique generative and adversarial learning strategy to leverage its own predictive ability. Finally, class activation visualization (Selvaraju et al., 2020) is used to understand how ESGAN exploits the information in the images to maximize its predictive ability.

## Results

### Benchmarks and ESGAN algorithms evaluation

As a baseline, all five model types were able to correctly classify whether plants have reached heading or not when provided with the full (100%) training data set of 3137 images (Figure 1, A, B, and H). The convolutional-based models CNN, ResNet-50 and ESGAN models all performed well (Overall accuracy [OA] = 0.89-0.92, F1-score = 0.87-0.90) and had superior performance than the tabular methods of KNN and RF (OA = 0.78-0.79, F1-score = 0.73-0.76).

**Figure 1.**
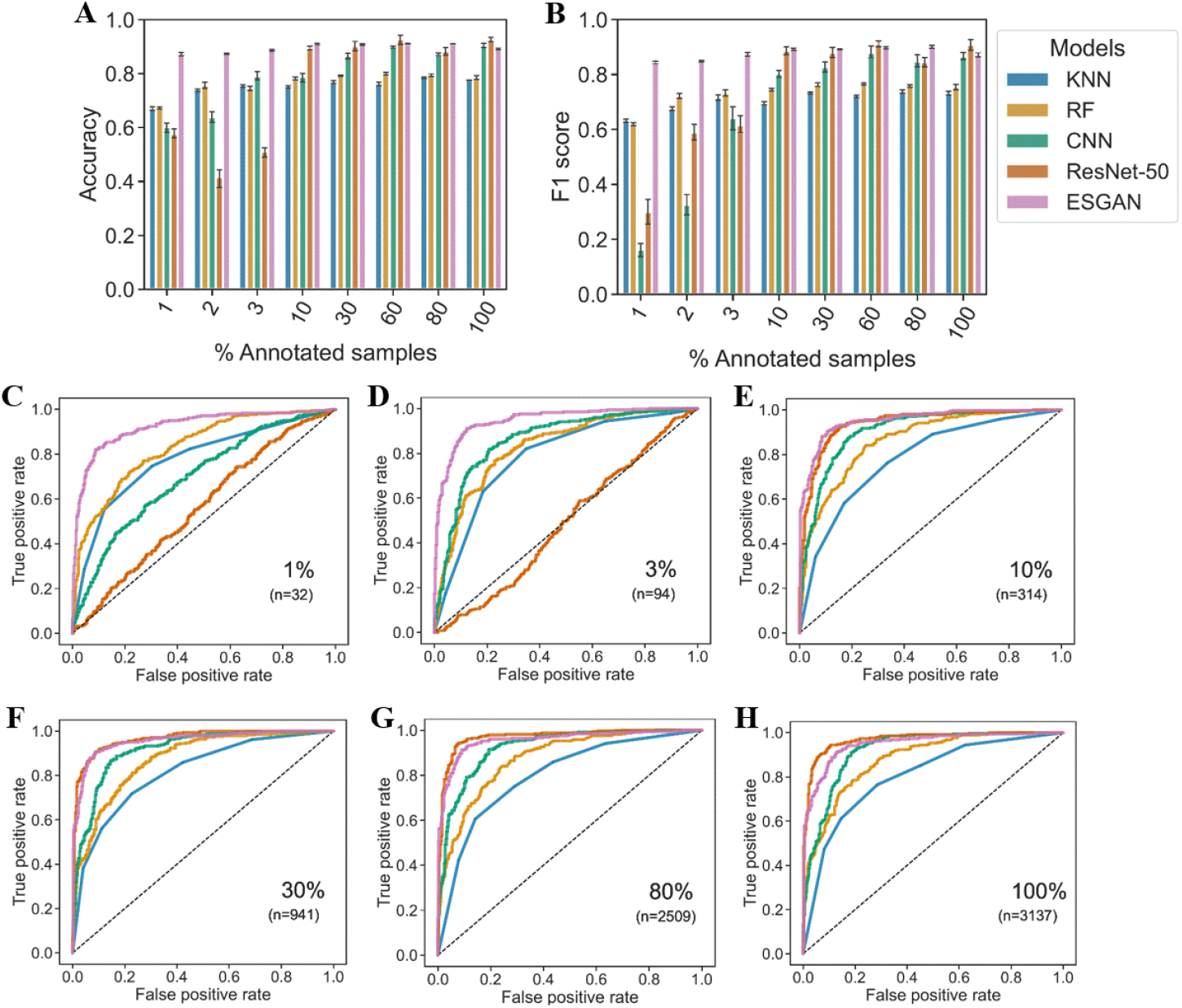
Evaluation of heading detection in testing data. Performance of benchmarks and ESGAN algorithms under an increasing number of annotated samples via Overall Accuracy (**A**) and F1-score (**B**) metrics. Error bars represent the standard deviation of performance metrics after 3 training and testing iterations. Performance evaluation using receiver operating characteristic (ROC) analysis is also presented for the same models under the same conditions (**C**-**H**). See Materials and Methods for explanation of metrics.

All model types demonstrated some reduction in ability to detect heading accurately as the amount of annotated training data was reduced, but to very different degrees. For ESGAN, the penalty for reducing the number annotated images used for training down to 1% of available data (32 images) was negligible in terms of OA (decline from 0.89 to 0.87), F1 score (decline from 0.87 to 0.85), and ROC analysis (Figure 1). TL using ResNet-50 was the next most robust method, maintaining performance as annotated training data was reduced to 10% (314 images), before being heavily penalized as the amount of annotated training data declined further (Figure 1). CNN performed at an intermediate level, maintaining performance as annotated training data was reduced to 30% (941 images), before being heavily penalized as the amount of annotated training data decline further (Figure 1). KNN and RF were less sensitive than CNN and ResNet-50 to reductions in the amount of annotated training data provided, but this only partially compensated for the poorer baseline performance of KNN and RF (Figure 1).

When the amount of annotated data was most restricted (1% of data available for training), ESGAN’s performance (OA = 0.87-0.89, F1 score = 0.85-0.87) was substantially better than all other models (OA = 0.43-0.75, F1 score = 0.16-0.72) (Figure 1, A and B). This also agreed with ROC analysis, where ESGAN was the most effective model for correctly identifying the two image classes when fewer than hundreds of annotated images were available for training (Figure 1, C and D).

### Understanding the predictive improvement of ESGAN

The ability of ESGAN to accurately determine heading of plants from aerial imagery can be explained by the synergic contributions of ESGAN’s generator and ESGAN’s discriminator sub models. The ability of the ESGAN generator to improve the visual representations of ‘fake’ images was notable during the training process (Figure 2, A, B, D, and E). The initial attempts of the ESGAN generator to generate images produced very noisy and unrealistic representations of *Miscanthus* plants (Figure 2, A and D). It was notable that the ESGAN generator sub model progressively learned to better match the RGB color intensity and spatial distribution of pixels of the real images turning them into very realistic representations of plants (Figure 2, B and E). This improvement was in agreement with the increasing performance of the ESGAN discriminator (Figure 2, C and F) along the successive minibatch steps of training, where the ability of this sub model to identify plants with panicles consistently improved regardless of whether very few (e.g. 32 images, Figure 2C) or many (Figure 2F) annotated training images were provided.

**Figure 2.**
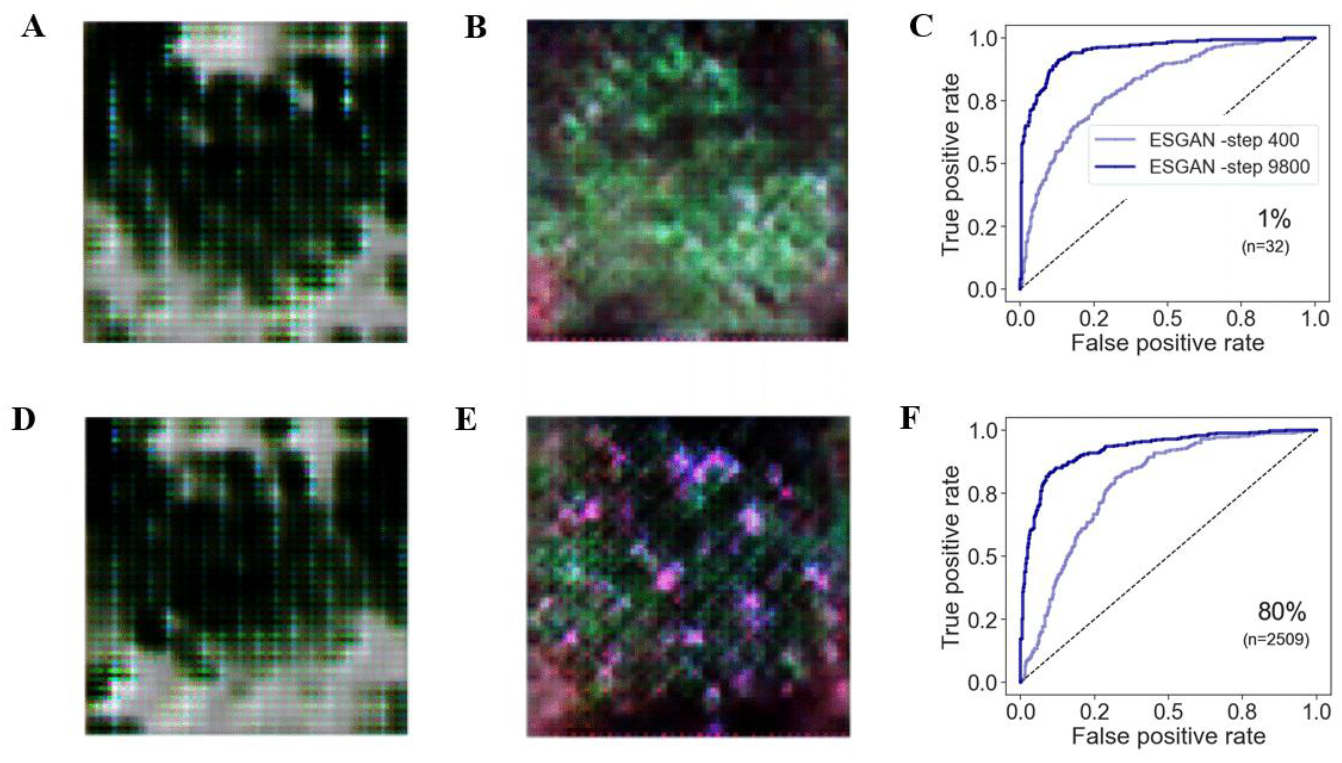
Visual representation of ‘fake’ images generated by the ESGAN generator during modeling implementation at early (400) (**A, D**) **and advanced** (9800) (**B, E**) **training steps**. Evaluation of heading detection by the ESGAN discriminator supervised classifier at early (400) (light blue) and advanced (9800) (blue) training steps under limited (1%) (**C**) and large (80%) (**F**) number of annotated samples.

### Explaining ESGAN’s learning via Grad-CAM

Since the ultimate goal of this study was to maximize the ability of the ESGAN discriminator supervised classifier to accurately determine the heading status of plants, gaining insights and interpretability about the learning process of this classifier was a key component of the analysis. When interpreting the learning process of this model via Gradient-weighted Class Activation Mapping (Grad-CAM) to highlight which parts of an image contribute the most to a model’s decision (Selvaraju et al., 2020), it was notable that the model successfully focused on plant pixels versus background pixels, and varied its activation levels depending on the class of image being considered. For plants without visible panicles (Figure 3A), higher activation regions (yellow) were visibly located over the green areas of the plant, this was especially notable over the upper leaves, while lower leaves and background regions (i.e., soil) were assigned with lower (blue) activation level (Figure 3C), meaning they were less informative. For the class of plant that had reached heading (Figure 3B), higher activation was particularly noticeable over the regions of panicles (i.e., silver-white objects) of the plants, while the model assigned lower activation level to vegetative tissues (Figure 3D).

**Figure 3.**
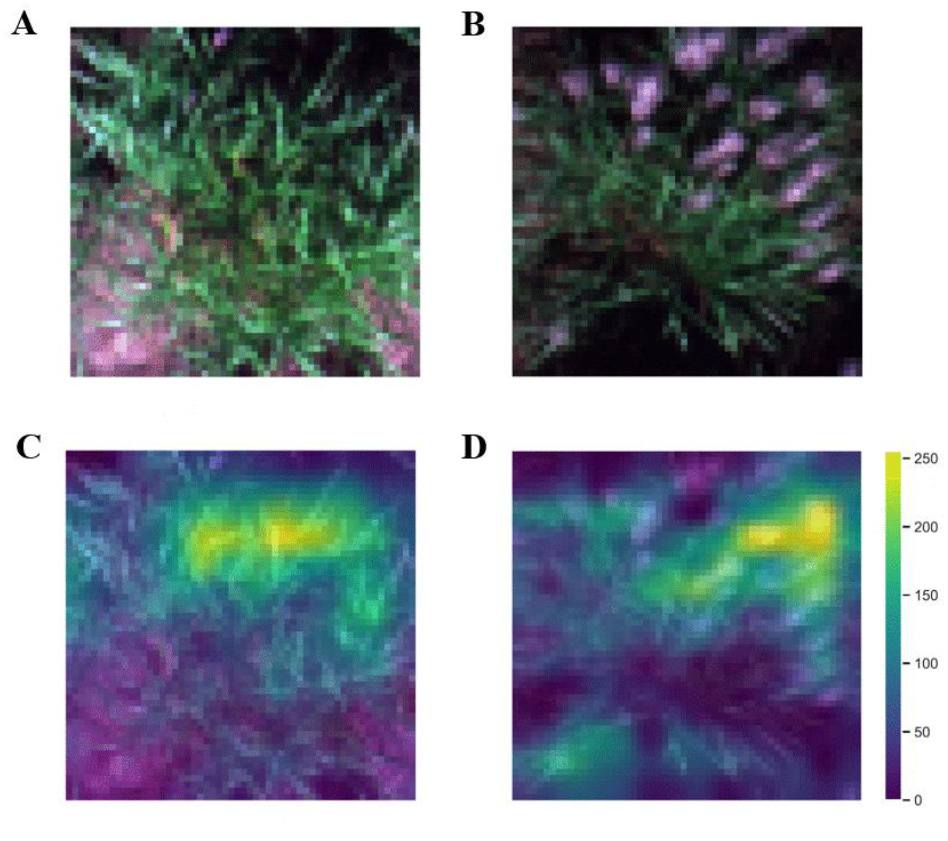
Visualization of examples real RGB images and Grad-CAM activation maps. Example pre-heading (A) plant class and the corresponding activation map (C) extracted from ESGAN D supervised classifier. Example post-heading (B) plant class and the corresponding activation map (D). The level of activation in the images are represented by low (blue), and high (yellow) values in a (0-255) scale.

### Evaluation of labor requirements for ESGAN versus human evaluation of heading status

The combined Miscanthus breeding trials studied here featured 3040 plots, including 12400 individual plants at the time of establishment (1 per plot for *M. sacchariflorus*, 10 per plot for *M. senensis*). Heading status of each plant was assessed on three occasions. Visual inspection by humans walking through the trials, including recording of data on an electronic device, required approximately 10.5 person-seconds per plant, or 36 person-hours in total on each occasion that phenotyping was performed (Table 1). By comparison, the time demand could be reduced >8-fold to 4.33 person-hours in total, or ∼1.2 seconds per plant, when acquiring images by UAV and analyzing them with ESGAN (Table 1). This reduction in time commitment reduces labor requirements below the threshold where, weather permitting, a single person could maximize the accuracy of heading data estimates by performing phenotyping on a daily basis.

**Table 1.** Description of activities and time required to phenotype the heading status of Miscanthus breeding trials by traditional visual inspection on the ground versus UAV imaging plus analysis by ESGAN. Data corresponds to the effort required to phenotype the three trials (3040 plots) in this study on one occasion i.e. at a single point in a seasonal time course.

| *Visual inspection by humans on the ground* |  | *UAV imaging and ESGAN analysis* |  |
| --- | --- | --- | --- |
| *Activity* | *Time* | *Activity* | *Time* |
| In-field evaluation and data recording on an electronic device. | 36 h | Flight planning and execution | 1 h 20 min |
|  |  | Image processing | 2 h 40 min |
|  |  | Image chip generation and ESGAN predictive inference | 20 min |
|  | **Total 36 h** |  | **Total 4 h 20 min** |

Before ESGAN can be deployed to analyze UAV imagery, it must be trained on human annotated images. The number of training images annotated by in-field, human phenotyping needed to maximize how accurately plants were classified as having reached heading or not, was substantially fewer for ESGAN (∼32 images) than for transfer learning by ResNet-50 (∼314) or a traditional, fully-supervised CNN (∼941 images). Based on the average time to phenotype each plant, this means that the time required to collect annotation data in each new context that a model would be used decreases by an order of magnitude for ESGAN relative to TL and CNN (Fig. 4a).

**Figure 4.**
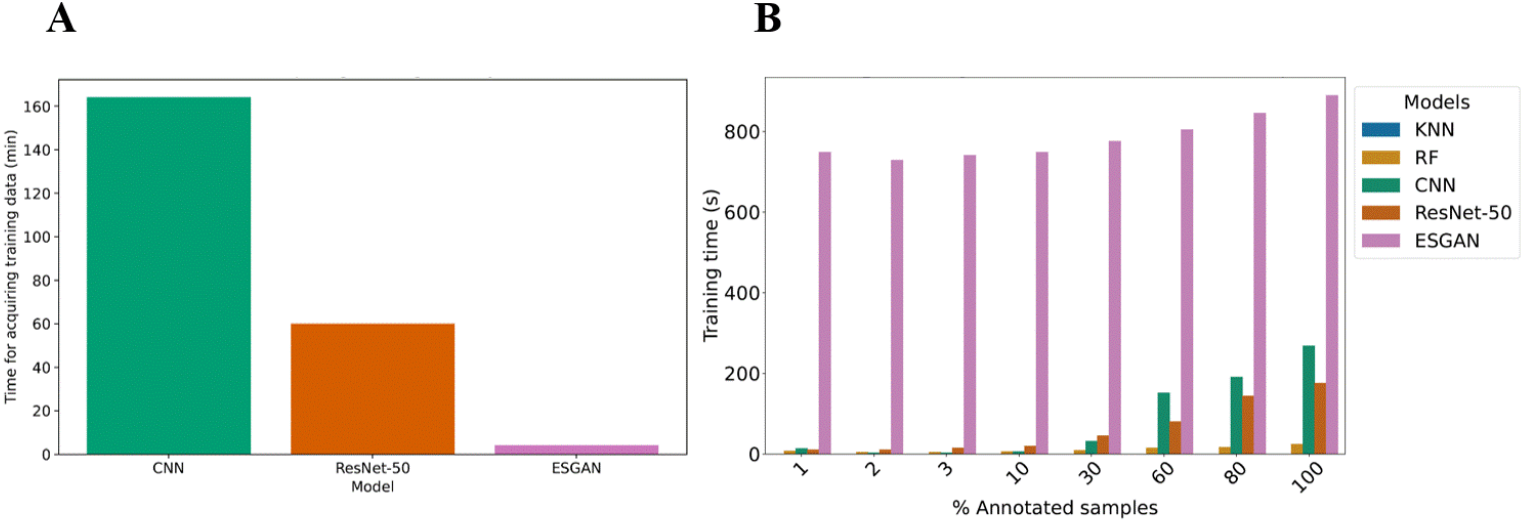
Evaluation of (A) time for acquiring annotation data for training for models that accurately classify images (OA > 0.85) and (B) training time for each model relative to the number of annotated samples in the training data.

In addition, the training time for ESGAN varied from ∼750-900 seconds depending on the number of annotated samples analyzed. This was 3-to 4-fold slower than for other learning methods (Figure 4B). However, this increase in computational time is small compared to the gains in efficiency with respect to field work (Figure 4A).

## Discussion

This study successfully demonstrated that a ESGAN approach can substantially reduce the amount of human-annotated training data needed to accurately perform an image classification task. Only tens of human-annotated images were needed to achieve high levels of accuracy in detecting plants that had reaching heading, or not, even when the problem was presented in the challenging context of a large population of *Miscanthus* genotypes, which feature a wide diversity of visual appearance both before and after heading. By contrast, hundreds of human-annotated images were needed to train a transfer learning tool (ResNet-50), and thousands of annotated images were needed to train a fully-supervised CNN. Meanwhile, KNN and RF were not able to classify images with high levels of accuracy, even when provided with thousands of training images. These findings highlight how a generative and adversarial learning strategy can provide an efficient solution to the common problem of needing large amounts of annotated training data for high-performing FSL DL approaches. This is a particularly significant discovery for the many potential applications of computer vision, such as high-throughput phenotyping in crop breeding, where frequent re-training of a DL model is needed to cope with strong context dependency of outcomes. The time required to acquire imagery by UAV and perform analysis with the ESGAN tool was ∼8-fold less than the time required for people to visually assess and record the heading status of Miscanthus while walking through the field trials. The time required to train any of the ML models is trivial relative to the time required for data acquisition. Combined with reducing the requirement for training data by 1-2 orders of magnitude by using ESGAN versus FSL or TL, this represents a major reduction in the effort needed to develop and use custom-trained ML models for phenotyping heading date in trials involving other locations, breeding populations, or species. For the Miscanthus breeding program at UIUC, the reduction in labor on each occasion the heading status of the breeding trials is assessed, from 36 person-hrs to 4.33 person-hrs, creates the opportunity to increase the frequency of assessment from once per week to once every two or three days, and thereby increase the accuracy of heading date estimates (Varela et al. 2022).

The power of ESGAN is valuable to research in the biological science domain, particularly at the intersection of remote sensing, precision agriculture, and plant breeding. The integration of automated data collection based on non-contact sensors and ESGAN can provide a cost-effective solution for exploiting large volumes of unannotated inputs, which can be collected at relatively low cost using remote sensing platforms. It can reduce dependence on large annotation data sets while achieving performance equivalent to traditional FSL approaches. Making these advances in a highly productive perennial grass, such as *Miscanthus*, is particularly important and challenging because these crops are more difficult to phenotype, i.e., highly segregating outbred populations with each individual genetically unique, and voluminous perennial plants that grow larger each year make field screening by humans on the ground more difficult and time-consuming than in annual, short-stature crops (Varela et al., 2022b). Implementing this ESGAN-enabled strategy may allow breeders to grow and evaluate larger populations in more locations as a means to accelerate crop improvement (Lewandowski et al., 2016) but at lower cost given the reduced dependence on manual annotation. It will be interesting to test how the ESGAN applied can be transferred to assess heading in other important crops including maize (*Zea mays*), sorghum (*Sorghum bicolor*), rice (*Oryza sativa*), wheat (*Triticum aestivum*) and switchgrass (*Panicum virgatum*), which also have panicles visible at the top of the canopy. The focus would shift to supplying a reduced number of highly quality and strategic annotations, while relying on the generative and adversarial element of the ESGAN to reduce the gap in predictive ability instead of depending on large data collection campaigns required for robust FSL implementations.

Previous studies have reported the use of high spatial resolution remote sensing imagery to detect reproductive organs in plants using pixel-wise classification (Kurtulmuş and Kavdir, 2014; Kumar et al., 2021) and morphological operations (Zhao et al., 2021). While these studies demonstrated relevant advances for rapid detection of maize tassels and sorghum panicles, they heavily relied on manual supervision-for example, to determine optimal features for the supervised classifier or morphological filtering. This makes these approaches more highly context dependent, where they would require continuous retraining when exposed to new cases (i.e., a different geographic location or year of the breeding trial in which environmental conditions alter crop appearance), which can present important challenges for successful scalability (Zhou et al., 2017). More recently, convolutional-based deep learning has been successfully implemented to detect reproductive organs of plants via classification (Zan et al., 2020) and object detection (Liu et al., 2020). The convolutional operation of the algorithm enables the model to fully exploit the information in the image, whereas both the intensity of the signal and the spatial arrangement of pixels in the image can be informative features to characterize the target trait (Yamashita et al., 2018). This also enables fully automatized feature extraction that provides a clear advantage over traditional machine learning, which heavily relies on expert-knowledge and manual feature engineering to identify meaningful features (O’Mahony et al., 2020). Nevertheless, the cost of creating a large volume of annotated datasets can be operationally and financially unfeasible in numerous applications (Weinstein, 2018; Lotter et al., 2021; Sager et al., 2021). Previous efforts to reduce dependency on training data have successfully integrated remote and proximal sensing with TL for plant species recognition (Letsoin et al., 2022), seedling detection (Tseng et al., 2022) and disease detection (Kamal et al., 2019). For example, the use of TL significantly reduced the computational time and resources needed when compared to implementing custom CNNs from scratch (Jha et al., 2019). While TL has proved an effective approach, the similarity between the original and target tasks can bring challenges for successful transferability; thus, additional data is often needed (Letsoin et al., 2022) to improve the generalization ability of TL. Moreover, the relationship between transferability, data annotation size, and predictive ability of TL has not been extensively tested in plant science applications. Therefore, the implementation of robust analytical approaches that directly address this challenge and accurately determine differences in the phenological characteristics of individuals in a population with diverse origins can be of significant interest for a challenging task that is typically done by manual inspection (Dong et al., 2021) and requires copious amount of manual annotation for robust FSL implementations (Varela et al., 2022a).

ESGAN clearly out-performed FSL models when only tens of training images were provided. Overall, this highlights the particular ability of ESGAN to exploit unannotated imagery to produce meaningful improvements for more accurate determination of the heading status under minimal annotation. This can be attributed to ESGAN’s ability to effectively enrich the latent space representation, which is crucial for classifiers in the discriminator to accurately distinguish between target classes and outperform other convolutional-based benchmark models. A previous study also used a GAN for discrimination of crops versus weeds (Khan et al., 2021) in high spatial resolution aerial imagery, but the reductions in demand for human-annotated training data were significantly less than achieved with ESGAN. This difference may reflect variations in the particular design of the ESGAN’s discriminator. ESGAN benefits from using two CNNs (supervised and non-supervised classifiers) that share weights, allowing synergic feature matching even when annotations are severely restricted. Specifically, the architecture design and training sequence of ESGAN ensure that weights updates in one classifiers affect the other one (Figure 7B), facilitating feature matching. This design and sequence of steps during training enable the model to synergistically exploit both types of data sources (i.e., annotated and unannotated), providing a clear advantage over the FSL and traditional TL strategies. Even though the main goal of this study was to maximize discriminator supervised classifier performance, the generative component of the algorithm showed a significant improvement in the quality of the visual representation of *Miscanthus* plants during the learning process (Figure 2A, B, D, and E). This allowed synergistic gains in the performance of the ESGAN discriminator and ESGAN generator as gradient updates and loss function information passed between sub models.

The dependence of the CNN model on voluminous amounts of annotated images was strong. This was not surprising and agreed with previous studies noting the importance of large and high quality datasets for optimal performance of FSL algorithms (Wang et al., 2021). This constraint was also evident, although to a lesser degree, when using the TL strategy. This demonstrated that the TL strategy was capable of exploiting prior knowledge, but the dependence on annotated images was consistently larger than for ESGAN.

The generally poorer performance of tabular-based algorithms (KNN, RF) versus CNN-based methods (ESGAN, TL, CNN) suggested that the use of spectral features summarized over an area of interest via statistical descriptors was a sub-optimal solution for the effective determination of heading status at the image level. This can in part be explained by the fact that inflorescences of the plants appeared as small silver color objects in the images, and it is logical to argue that the use of spectral descriptors extracted and summarized over an area of interest (i.e., whole image chip) into a unique tabular value may only partially capture these patterns in the image. The capacity of convolutional-based approaches to automatically map spatial-dependent patterns in the images and use them as informative features (Wei et al., 2019) can explain the superior ability of convolutional-based approaches.

Grad-CAM showed that the algorithm prioritized information gain from areas of the image occupied by inflorescences and vegetative tissue as a means to differentiate each class without the need for manual supervision to identify regions of interest. This extends the degree to which expert supervision was not needed during implementation of the analysis. This is particularly important in biological systems, such as crop breeding, where high levels of phenotypic diversity from genetic and environmental sources occur, which would otherwise limit the broad application of existing AI tools.

By reducing the dependence from manual annotation, the traditional requirement for exhaustive field-wide surveying can be alleviated also to determine the heading dynamics. This has important implications for optimizing the operational costs associated with phenotyping trials. Rather than conducting comprehensive surveys of the entire field at each round of evaluations, surveying could focus on representative sections to optimize the operational cost. Complementing these targeted ground surveys with aerial surveys would further enhance temporal coverage by better distributing the operational cost associated, without compromising accuracy in heading status predictions and reduce the cost of capturing finer temporal dynamics. ESGAN’s strong predictive performance even with reduced data availability suggests that this hybrid approach could maintain high levels of accuracy across time points, offering a practical, cost-efficient, and scalable alternative for large-scale phenotyping in agricultural research.

## Conclusion

The generative-discriminative nature of the proposed ESGAN approach, which is characterized by an adversarial and synergic training of neural networks presents a promising avenue for reducing the dependence on human supervision and enhancing the model’s capacity for generalization through more efficient incorporation of contextual information. In the case study presented, this meant that heading detection in plants could be effectively determined using high-spatial-resolution aerial imagery with only very limited (tens) of human-annotated training images. This represents a significant potential reduction of manual annotation given by ESGAN compared to traditional FSL, all with negligible penalization. These outcomes are valuable for designing future strategies to optimize the integration of manual field screening efforts and aerial data collection. More broadly, this work could address the need for advanced modeling techniques that can produce both robust accuracy while reducing the operational cost of collecting time-consuming annotated data for many computer vision problems in plant science applications.

## Materials and Methods

### Field trials

Data was collected from three *Miscanthus* diversity trials located at the University of Illinois Energy Farm, Urbana (40.06722°N, 88.19583°W). The trials were planted in the spring of 2019. This study focused on the second year (2020) of their establishment, which is the first growing season in which the *Miscanthus* trials are typically phenotyped. The broader aims of the breeding program include assessment of overwintering survival and evaluation of germplasm adapted to a wide range of latitudes and environments. Since plants that were lost to lethal winter temperatures were randomly distributed within the trials, all locations were phenotyped by humans and UAV imaging regardless of survival status. Not all germplasm experienced the photoperiod necessary to achieve a vegetative-reproductive transition and achieve heading at this location.

The *M. sacchariflorus* trial included 2000 entries as single-plant plots in four blocks, each block including 58 genetic backgrounds (half-sib families) (Varela et al., 2022b). The size of the trial was 79 m long × 97 m wide, and each plot (plant) was 1.83 × 1.83 m size.

One *M. sinensis* trial included germplasm from South Japan while the other included germplasm from Central Japan (Clark et al., 2015). Each of these two trials included two blocks, with 130 plots per block. Each plot contained seedlings from a single half-sib family, with 10 plants at a spacing of 0.91 m, requiring transplant of 10,400 individuals in total at the start of the trial. In the *M. sinensis* South Japan trial there were 124 families, and in the Central Japan trial there were 117 families. Therefore, a few families were planted in more than one plot per block to avoid leaving empty space. The size of the field that included both of the *M. sinensis* trials was 115 m long × 121 m wide.

### Trait of Interest and Ground-Truthing

Every plant in both single-plant and multi-plant plots was phenotyped individually through observation on the ground by an expert evaluator to determine if it had produced panicles or not. A plant was considered to have reached heading once the culms that contribute to the canopy height have 50 percent panicles that had emerged ≥ 1 cm beyond the flag leaf sheath. Data were recorded to separately track plants that died or never reached heading. Examples of plants with emerging panicles imaged at the ground level and by UAV are shown in Figure 5. The *M. sacchariflorus* trial was inspected on DOYs 248, 262, 276, and the *M. sinensis* on DOYs 245, 265, and 280. This matched as close as possible (i.e., depending on optimal weather conditions) the dates of UAV data collection.

**Figure 5.**
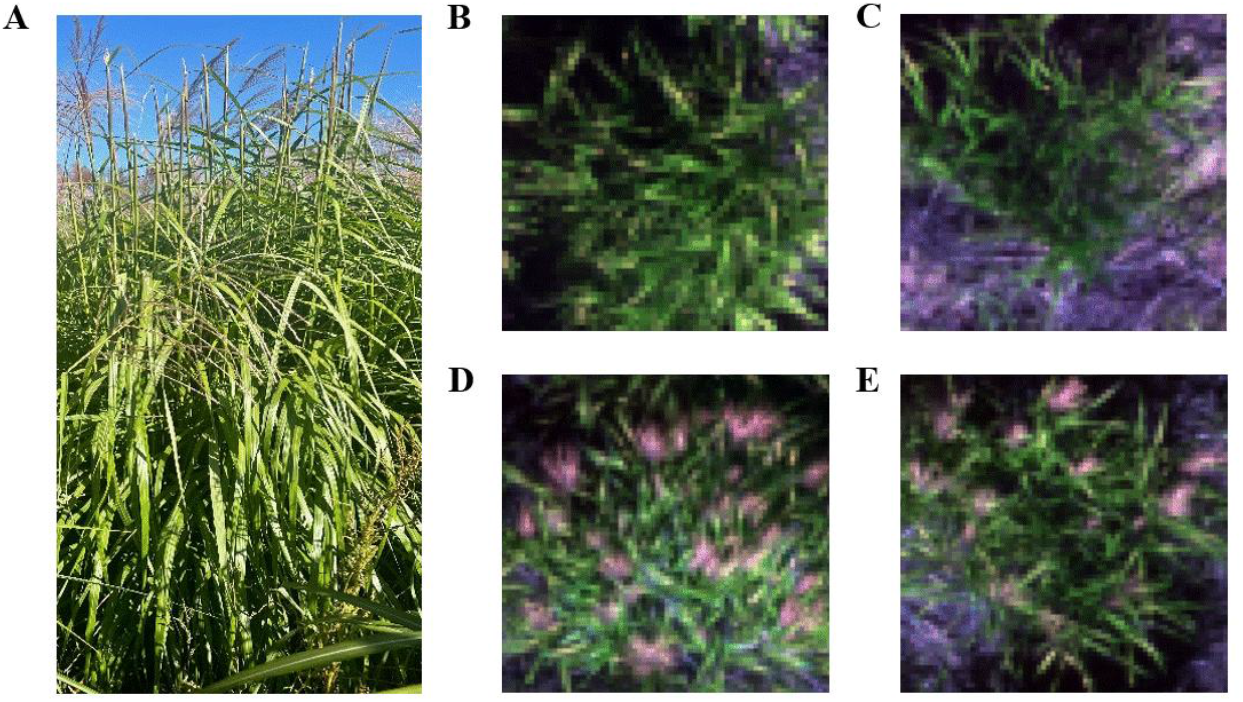
Example cases of plant with emerging inflorescences from ground (A), plants not yet heading (B, C) and plants after heading (D, E) from UAV imagery collected in the 2020 season.

### Aerial Data Acquisition and Imagery Preprocessing

A Matrice 600 Pro hexacopter (DJI, Shenzhen, China) UAV equipped with a Gremsy T1 gimbal (Gremsy, Ho Chi Minh, Vietnam) mounted with a multispectral RedEdge-M sensor (Micasense, Seattle, WA, USA) was utilized for aerial data collection. The sensor included five spectral bands in the blue (465 to 485 nm), green (550 to 570 nm), red (663 to 673 nm), rededge (712 to 722 nm), and near-infrared (820 to 860 nm) regions of the electromagnetic spectrum. Flights were conducted 3 times (DOYs 247, 262, and 279) in the season corresponding to the period when most inflorescences emerge. The aerial data was collected under clear sky conditions around ±1 hour from solar noon to ensure consistent reflectance values across days of data collection. The flight altitude was 20 m above ground level, resulting in a ground sampling distance (GSD) of 0.8 cm/pixel. Flight settings included 90% forward and 80% side overlapping during data acquisition to ensure high quality image stitching during post processing steps. Ten black and white square panels (70 cm x 70 cm) were distributed in the trials as ground control points (GCPs). An RTK (real-time kinematic) survey was done using a Trimble R8 global navigation satellite system (GNSS) integrated with CORS-ILUC local station to survey the GCPs to ensure consistent spatial extraction of the images chips between days of data collection. A Micasense calibration panel was imaged on the ground before and after each of the flights for spectral calibration of the images via an empirical procedure (Poncet et al., 2019). Images were imported into Metashape version 1.7.4 (Agisoft, St. Petersburg, Russia) to generate calibrated surface reflectance multispectral orthophotos. Image processing and analysis was performed with a i9-12900H processor, with 14 cores 32GB RAM, and a NVIDIA GeForce RTX 3080 16GB GPU. The orthophotos from each of the three sampling dates were resampled to a common 0.8 cm/pixel resolution and stacked into a 3-band RGB (i.e., red, green, blue bands) raster stack object. Further steps in the analysis consider only the RGB bands of the multispectral sensor for the following reasons: 1) RGB has proved to be highly sensitive and competitive with the red edge and near infrared spectral regions of the electromagnetic spectrum for monitoring heading in *Miscanthus* (Varela et al., 2022b). 2) The use of RGB bands allowed testing of TL as potential alternative approach into the analysis. Image chips for each plot/plant were generated by clipping the stacked orthophotos objects using a polygonal shapefile that includes each plot polygon of the trials. The resulting image chips containing the 3-dates of RGB bands were further split into single date matrix arrays in Python for further analysis. The size of the images chips was 108 pixels × 108 pixels × 3 RGB bands per date.

### Dataset

After accounting for plants that died due to lethal winter temperatures or never reached heading, a subset of 1309 genetically diverse plants were identified for which ground truth data and UAV imagery were available on each of the three sampling dates during the growing season. This resulted in a dataset of 3921 instances of single-plant images and associated heading status.

### Algorithms

#### KNN and RF

KNN is an extensively used algorithm for pattern classification (Weinberger et al., 2005). The proximity distance between individuals is used to determine class discrimination in a population. The core concept is that the closer the individuals are in the feature space, the higher probability of belonging to the same class (Short and Fukunaga, 1981). The advantage of this method is the reduced number of parameters and fast computation, while the down side are the sensitivity to irrelevant features and difficulty for determining the optimal value of the parameter number of neighbors. In our study, after preliminary experimentation, parameter number of neighbor was set equal to 10.

RF is a versatile non-parametric algorithm that has been broadly used in classification tasks (Belgiu and Drăguţ, 2016). It exploits bagging and feature randomness to build an ensemble of trees in which prediction by committee tends to be more accurate than in any of the individual trees (Breiman, 2001). RF is straightforward to use and requires simple hyper-parameter tuning to deliver high predictive performance. Another advantage of this algorithm is that it does not assume normal distribution of data or any form of association between the predictors and the response variable (Probst et al., 2019). Furthermore, as an ensemble of trees, RF is highly capable for managing overfitting. For implementation, parameters number of estimators and maximum depth of trees were optimized via *GridSearchCV* function in Python.

The KNN and RF algorithms require tabular features as inputs for modeling implementation. Tabular-based features were generated from the image chips using *Numpy* Stats functions in Python. Statistical descriptors median, range, standard deviation, percentile 75, percentile 95, and percentile 99 values were utilized to extract tabular features values from the RGB bands of each of the image chips (based on structural and multispectral bands not contributing additional explanatory power in prior assessment of heading by Miscanthus in UAV images, Varela et al. 2022). This process generated a total number of 18 features that were further used as inputs of the algorithms to determine the heading status of each of the plants (i.e., image chip level).

#### Custom CNN

CNN is a deep learning technique successfully utilized for image analysis (Teuwen and Moriakov, 2020). The architecture of the algorithm consists of a series of hidden layers that map the input images to output values (Yamashita et al., 2018). The core component of the algorithm is the convolution operation, where a set of trainable kernels are applied to the input image to automatically generate a set of spatial features that best describe the target predictor. The model learns basic features in the first layers, and more complex feature representations at deeper layers iteratively (i.e., via gradient loss and backpropagation) (Szegedy et al., 2015). The typical architecture of the algorithm includes a backbone feature generator and classifier or regressor head. In this study, the backbone feature extractor of the custom CNN includes six convolutional layers all including maximum-pooling and batch normalization. Convolutional layers 2, 4, and 6 additionally consider 40% features dropout, flattening layer. Then the backbone feature extractor also included a fully connected layer, batch normalization, and 50% feature dropout. Finally, the classifier head of the CNN includes a sigmoid activation layer that delivers predictions as normalized probability distribution values (i.e., with panicles or without panicles) for each image chip. After preliminary experimentation, the number of features in each layer was set to 32, 32, 64, 64, 128, and 128. Zero padding, stride equal to one with no overlapping, and ReLU (rectified linear unit) activation function were also considered in the architecture design. The CNN’s kernel filter size was set to 3 pixels × 3 pixels × 3 RGB bands, and max-pooling was set equal to 2 following each convolution. Binary cross entropy was utilized as loss function of the classifier head of the neural network.

#### ResNet-50

TL is a deep learning technique that exploits stored knowledge gained while solving one problem that can be then applied to solve a different but related task (Yang, 2008). This prior knowledge is stored in large neural networks, and then transferred to solve a target task (Ribani and Marengoni, 2019). This implies several advantages over training a custom CNN from scratch, for e.g., reduction in computational resources and latency for delivering predictions (Jha et al., 2019), and boost in predictive ability over the target task (Wang et al., 2018). Deep neural networks trained on ImageNet dataset have reported state-of-the-art performance in TL applications (Kornblith et al., 2019). ResNet-50 (Deng et al., 2009) is a deep-50 layers neural network specifically designed to exploit residual connections between convolutional layers trained on large ImageNet dataset. This ensures that weights learned from previous layers do not vanish during back propagation (Noh, 2021), which represents an advantageous trick in the design that enables the use of a large number (i.e., deeper) of layers in the architecture of the network. For implementation, we followed the steps suggested by (Kornblith et al., 2019), where the strategy is as follows: 1) remove the original head of the pretrained neural network, 2) add a custom binary classifier head, 3) fine-tune the top 5 layers while keeping bottom layers frozen. ResNet-50’s pretrained weights and biases were imported from Keras (Chollet, 2015a). The original image chips were resampled to a 128-pixel size to fit the input image size of the ResNet-50 network.

#### ESGAN

The core concept of GAN involves training deep generative networks based on game theory (*62*) (Goodfellow et al., 2014). The model contains two CNN sub models: 1) a generator (G), and 2) a discriminator (D) that are trained in an adversarial manner. Both G and D are trained to optimize the results, where the goal of G is to mislead D, and the goal of D is to distinguish between fake images generated by G and real images collected with the UAV. GAN has been successfully implemented for image generation (Ahmad et al., 2022), augmentation (Sandfort et al., 2019), and classification (He et al., 2017) tasks. Adapting GANs to the semi-supervised context by forcing the discriminator network to output class labels was discussed in (Odena, 2016) (Erickson et al., 2017). We explore this direction in the context of plant science applications. During the training process, the data generated by the generator (G) is used to train the discriminator (D). This process enabled D not only distinguish between real and fake data but also to identify whether a plant has reached the heading stage (Figure 6). Therefore, it is expected that D can learn features that allow for the discrimination of images with plants prior to, or after, heading using much less human annotated training data than in FSL.

**Figure 6.**
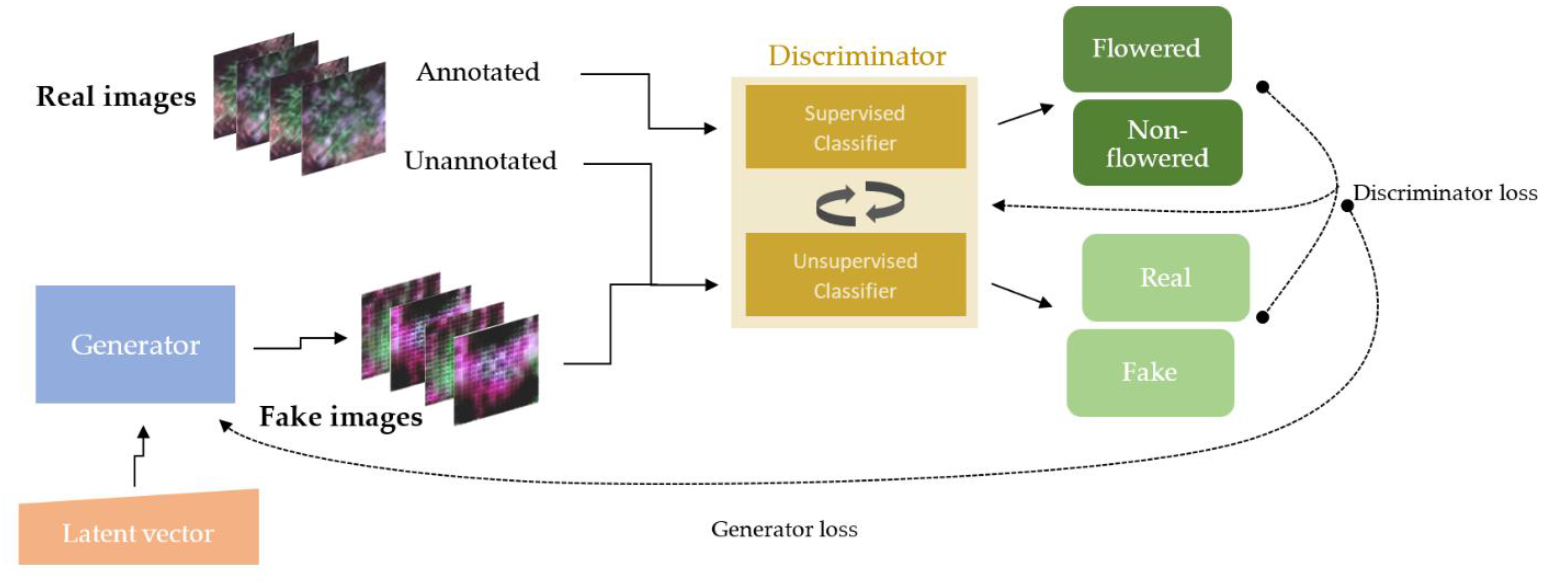
Diagram of ESGAN and data workflow including the generator (G) and discriminator (D) sub models utilized to assess flowering status.

**Figure 7.**
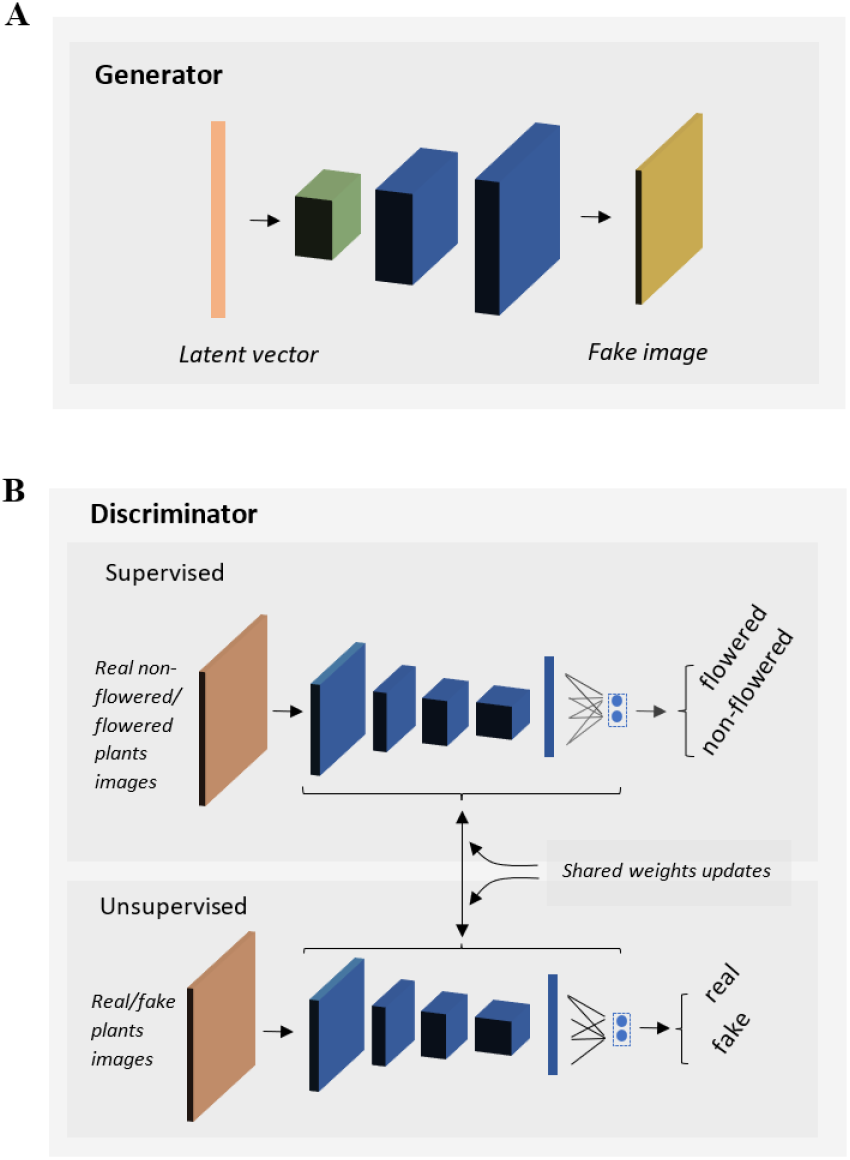
Diagram of ESGAN architecture. Components include: G (**A**) and D (**B**) sub models with the corresponding inputs (orange vector and cuboid), hidden layers (green, and blue cuboids and vectors), and outputs (i.e., yellow cuboid as fake image in G and classes predictions in D).

ESGAN was implemented following the steps suggested in (Salimans et al., 2016), while creating separated classifiers for supervised and unsupervised D. First, the D supervised classifier is implemented to infer the two classes (i.e., plants prior to heading or after heading) from real images using Softmax activation function. Then the supervised D produces predictive outputs for each image (i.e., between 0-1), which represent the normalized probability of the image belonging to the two image classes. The D unsupervised classifier is implemented by taking D supervised prior to Softmax activation (i.e., D supervised backbone feature extractor) re-using its feature extraction layers weights. It then calculates the normalized sum of exponential outputs (i.e., between 0-1) via a custom function, which represents the probability of the image being real or fake. This means that updates to one of the classifier models will impact both models.

The supervised loss function (*L* _*D supervised*_) is defined as the negative log probability of y when the correct class is allocated by x. *L* _*D supervised*_ focuses on correctly classifying input images to given labels.

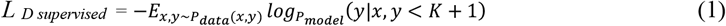

Unlabeled image loss functions constitute the unsupervised loss function (*L* _*D unsupervised*_).

≥(y = K+1|x) represents the probability that x is fake (i.e., traditional GAN), corresponding to 1-D(x) of GAN architecture. *X*_*u*_ denotes unannotated data samples. The unannotated real images are classified to one of the K classes by the first term of *L* _*D unsupervised*_. The second term in the *L* _*D unsupervised*_ classifies the images generated by the G as K+1 (fake).

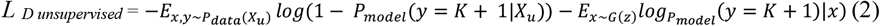

By minimizing *L* _*D supervised*_ and *L* _*D unsupervised*_, the classifiers are trained with gradient descent. D weights are stochastically updated by their gradient (Eq 3.) at each training step via gradient descent of Equations 1 and 2.

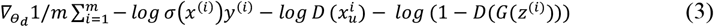

For all m samples in a minibatch σ(x)j = ≥(y = j|x) (SoftMax function) was applied at the output of D supervised. After some preliminary experimentation, a 72-pixel image size was used as inputs for CNN and ESGAN given the negligent penalization in predictive performance but significant saving on computational time. The G and D’s architectures (Figure 7).

Balanced sampling between annotated and unannotated images at each minibatch iteration was considered to ensure consistent performance of ESGAN during training. G initializes with a latent vector (Figure 7A, orange vector) as input, which is then reshaped (Figure 7A, green cuboid) and upscaled through two deconvolution (i.e., transpose convolution) operations (Figure 6A, blue cuboids) into a fake image (Figure 7A, yellow cuboid) that ensures match to the size of real images (Figure 7A, yellow cuboid) as output of G. D inputs both real and fake 72 × 72 × 3 (Figure 7B, orange cuboid) images. It is then followed by four convolutional operations and Max-Pooling layers of size two, followed by a flattening layer and 40% features drop out. The size of the convolutional kernel was 3 × 3 and the Leaky ReLU activation function was applied to all the layers of G and D, except for the output of G, which used the Tanh function. The standard Adam optimizer and learning rate equal to 0.0001 was employed in G and D sub models. The size of the convolutional kernel was 3 × 3. Each classifier could predict the input data to a label y from two K classes (plants with or without panicles) or to a fake sample (k+1 class).

### Algorithm Implementation and Metrics

KNN and RF were implemented using Scikit-learn library, while CNN, ResNet-50, and ESGAN were implemented in Keras, both in Python version 3.9.16. Each model fitting was iterated 3 times using a random training and testing partition to ease the convergence of the models’ prediction metrics. The number of image chips with the corresponding ground-truth data was 3921. 2021 images came from the *M. Sacchariflorus* trial and 1900 came from the *M. Sinensis* trials. The full dataset was split (80:20) into training and testing datasets. The training dataset was split further (80:20) into training and validation datasets. The validation dataset was used to optimize the models’ performance and prevent overfitting during training. CNN and ResNet-50 were trained for up to 300 epochs, while ESGAN was trained for up to 1000 epochs. Early stopping strategies were incorporated in these three models to prevent overfitting, optimize performance, and reduce computational time. The test dataset was utilized to expose the models to unseen data to evaluate the generalization ability of the models. As the number of annotated images used for training was altered to generate the eight sample size cases (Figure 1, A and B), the number of test images was held constant at the equivalent of 20% of the full dataset. This ensures that all models are evaluated on the same test set and size. Training was implemented on batches

Description of batch training loop of ESGAN:

1. train D with frozen G weights (i.e., non-trainable):
  a. train supervised classifier with annotated images, determine *L* _*Dsupervised*,_ gradient computation, and D weights update.
  b. train unsupervised classifier with real images, determine *L* _*Dunsupervised*_, gradient computation, and D weights update.
  c. train unsupervised classifier with fake images, determine *L* _*Dunsupervised*_, gradient computation, and D weights update.
2. train G with frozen D weights (i.e., non-trainable):
  a. feed D with G fake images.
  b. extract features and calculate G loss.
  c. update G weights by their gradient.

The overall accuracy (OA), F1 score, and ROC curve analysis were utilized as performance metrics of the models on classifying heading status (i.e. with or without visible panicles). OA and F1 score metrics are described in Equations 5-6:

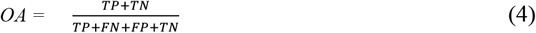

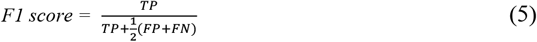

In Equations (5)–(6), true positive (TP) is defined as plants with panicles correctly classified as plants with panicles. True negative (TN) is defined as plants without panicles correctly classified as plants without panicles. False positive (FP) is defined as plants without panicles (i.e., ground-truth) incorrectly classified as plants with panicles (i.e., positive class). False negative (FN) is defined as plants with panicles (i.e., ground-truth) incorrectly classified as plants without panicles (i.e., negative class).

Receiver operating characteristic (ROC) analysis is especially useful for assessing models where the output is a probability score that can be thresholded to produce binary decisions. The technique involves plotting the ROC curve, which is a graphical representation of a classifier’s diagnostic ability between TP rate and FP rate at various threshold settings. The area under the ROC curve quantifies the overall ability of the classifier to discriminate between positive and negatives classes (Fawcett, 2006).

Grad-CAM is a technique used in deep learning to visualize which parts of an image contribute the most to a model’s decision. It highlights the regions of an input image that were more important for making a specific prediction (Selvaraju et al., 2020). The technique was implemented to improve the interpretation of the ESGAN D supervised classifier’s learning process. The visualizing technique highlights the importance of different regions of the image in the output prediction by projecting back the weights of the output layer on to the convolutional feature maps (Zhou et al., 2015). As recommended by (Chollet, 2015b), the following steps were used to generate the class activation maps. First, the ESGAN D supervised classifier mapped the input image to the activations of the last convolution layer as well as the output predictions. The gradient of the predicted value for the input image with respect to the activations of the last convolution layer was computed. Each image channel in the feature map array was weighed by how important this channel was with regard to the predicted value, and then all the channels were summed to generate the corresponding activation map array. The Grad-CAM activation map provided a measure of how strongly portions of the image contributed to the predictions made by the ESGAN D supervised classifier visualized in a 0-255 scale map array.

## Acknowledgments

We thank Tim Mies at the Energy Farm at University of Illinois for technical assistance.

## Author Contributions

S.V. and A.D.B.L. conceived the study, interpreted the data, and wrote the manuscript. E.S, X.Z, and J.N established, maintained, and collected the ground-truthing data in the field trials. J.R and D.A collected the aerial data. S.V implemented the data pipeline in the study.

## Data and code availability

The datasets used and coding implementation during the current study are publicly available from the online Illinois Databank at https://doi.org/10.13012/B2IDB-8462244_V2 and GitHub repository https://github.com/pixelvar79/ESGAN-Flowering-Detection-paper.

